# Structural properties of the HNF-1A transactivation domain

**DOI:** 10.1101/2023.06.23.546236

**Authors:** Laura Kind, Mark Driver, Arne Raasakka, Patrick R. Onck, Pål Rasmus Njølstad, Thomas Arnesen, Petri Kursula

## Abstract

Hepatocyte nuclear factor 1α (HNF-1A) is a transcription factor with important gene regulatory roles in pancreatic β-cells. *HNF1A* gene variants are associated with a monogenic form of diabetes (HNF1A-MODY) or an increased risk for type 2 diabetes. While several pancreatic target genes of HNF-1A have been described, a lack of knowledge regarding the structure-function relationships in HNF-1A prohibits a detailed understanding of HNF-1A-mediated gene transcription, which is important for precision medicine and improved patient care. Therefore, we aimed to characterize the understudied transactivation domain (TAD) of HNF-1A *in vitro*. We present a bioinformatic approach to dissect the TAD sequence, analyzing protein structure, sequence composition, sequence conservation, and the existence of protein interaction motifs. Moreover, we developed the first protocol for the recombinant expression and purification of the HNF-1A TAD. Small-angle X-ray scattering and synchrotron radiation circular dichroism suggested a disordered conformation for the TAD. Furthermore, we present functional data on HNF-1A undergoing liquid-liquid phase separation, which is in line with *in silico* predictions and may be of biological relevance for gene transcriptional processes in pancreatic β-cells.

## Introduction

Hepatocyte nuclear factor (HNF-1A) is a transcription factor with essential gene regulatory functions in the pancreatic β-cells, which are important regulators of glucose homeostasis. The β-cell-specific signaling cascade ‘glucose-stimulated insulin secretion’ (GSIS) is initiated by a rise in blood glucose levels after the ingestion of a meal and results in the secretion of insulin into the blood stream (1), in turn stimulating peripheral tissues to internalize glucose from the blood. HNF-1A is indispensable for β-cell maintenance and function, as it transcriptionally regulates components of the GSIS cascade, as well as various transcription factors integrated into a gene regulatory network (2-5). Understanding HNF-1A function is potentially clinically relevant, as numerous genetic variants within the *HNF1A* gene can cause the hereditary diabetes type ‘maturity onset diabetes of the young’ (HNF1A-MODY), be associated with HNF1A-MODY with reduced penetrance, or act as polygenic risk factors for type 2 diabetes (6-9).

HNF-1A exhibits a multi-domain architecture typical for transcription factors (Fig. 1A). The N-terminal dimerization domain (DD, residues 1 – 33) contains a helix-turn-helix motif and mediates homodimerization of HNF-1A or heterodimerization with the homologous protein HNF-1B (10-13). The central α-helical DNA-binding domain (DBD, residues 83 – 279) is composed of a POU-specific domain (POU_S_) and a POU homeodomain (POU_H_), which together bind to the promoters of HNF-1A target genes (14). The C-terminal region of HNF-1A constitutes the transactivation domain (TAD, residues 280 – 631), which has remained undercharacterized. Three HNF-1A isoforms have been identified, which differ in the length of the TAD and exhibit varying levels of gene transactivation potentials (15).

**Fig. 1.**
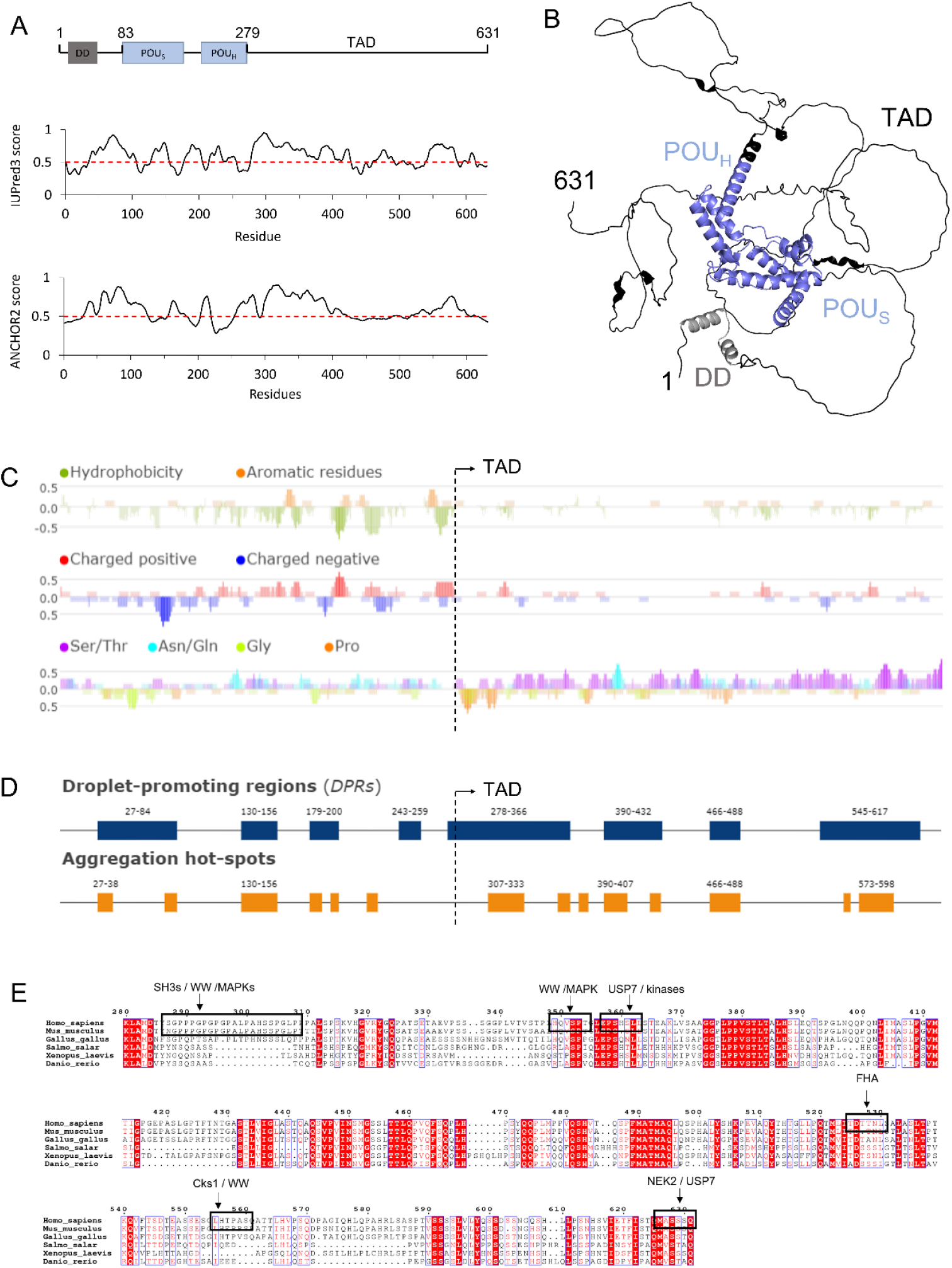
Structure prediction and sequence analysis of HNF-1A. **A**. Top: Domain overview of full-length HNF-1A with residue numbers indicated above. DD – Dimerization domain. POU^S^ – POU-specific domain. POU^H^ – POU homeodomain. TAD – Transactivation domain. Middle: IUPred3 (23) disorder prediction for full-length HNF-1A. Values above the threshold (red dashed line) indicate tendency for disorder, while values below the threshold indicate tendency for order. Bottom: ANCHOR2 (24) prediction of protein regions that undergo disorder-to-order transitions upon binding. High values represent highly disordered binding regions. **B**. AlphaFold model for full-length HNF-1A (26, 27). Coloring based on the domain presentation in A. **C**. Fast Estimator of Latent Local Structure (FELLS) sequence analysis for full-length HNF-1A regarding residue hydrophobicity (top), amino acid charge (middle), and compositional bias towards Ser/Thr, Asn/Gln, Gly, and Pro residues (bottom). **D**. FuzDrop predictions of droplet-promoting regions and aggregation hotspots in full-length HNF-1A. **E**. Multiple sequence alignment of HNF-1A TAD (residues 280-631) for the model organisms *Homo sapiens, Mus musculus, Gallus gallus, Salmo salar, Xenopus laevis, Danio rerio*. Conserved residues are marked with red background, and similar residues are denoted in red notation. SLiMs retrieved from the ELM database are depicted by a black box and specified by the description above.

Eukaryotic transcription factors commonly contain regulatory domains (16, 17), which can alter gene transcription in an activating or inhibiting manner, e.g. by binding to components of the transcriptional machinery or by mediating chromatin remodeling (18). These regulatory domains frequently contain intrinsically disordered regions (IDRs), which often exhibit a compositional sequence bias (16, 19). HNF-1A likely harbors such IDRs and several sites for protein-protein interactions in the C-terminal TAD. Specific regions within the TAD have been shown to promote transcriptional activity of HNF-1A (residue ranges 398-470, 544-631, and 440-506) and are thus proposed to function as activation domains (20, 21). However, detailed knowledge on the structural and functional features of the HNF-1A TAD is currently lacking, preventing a mechanistic understanding of the gene transcriptional functions of HNF-1A and the impact of *HNF1A* variants within this region.

Here, we present a bioinformatic approach to identify features in the TAD of HNF-1A that may be of functional importance in transcriptional regulation by HNF-1A. We further developed a protocol to recombinantly express and purify a TAD-containing HNF-1A protein and assessed the behavior of the TAD in solution. Finally, we present functional data supporting a liquid-liquid phase separation (LLPS) behavior of HNF-1A, which may play a role in gene transcriptional processes of pancreatic β-cells. This knowledge is important for the development of novel approaches in precision medicine and can lead to improved care for patients with HNF1A-MODY.

## Materials and Methods

### Bioinformatic analyses

HNF-1A protein sequences from various model organisms were extracted from the UniProt (22) database under the following accession codes: *Homo sapiens* (human): P20823, *Mus musculus* (mouse): P22361, *Gallus gallus* (chicken): Q90867, *Salmo salar* (Atlantic salmon): Q91474, *Xenopus laevis* (African clawed frog): Q05041, *Danio rerio* (zebrafish): Q8UVH4.

Disorder predictions of full-length *Hs*HNF-1A were performed by using the IUPred3 (23), ANCHOR2 (24) and the Fast Estimator of Latent Local Structure (FELLS) (25) webservers. The FELLS output was further utilized to investigate sequence composition, such as residue hydrophobicity, charge, and sequence compositional bias. AlphaFold (26, 27) was employed for a tertiary structure prediction of full-length *Hs*HNF-1A. The FuzDrop algorithm (28) was used to predict the tendency of *Hs*HNF-1A to undergo LLPS. A multiple sequence alignment of the HNF-1A TAD (*Hs*HNF-1A, residues 280-631) was generated using the Clustal Omega alignment tool (29, 30) and visualized using ESPript3.0 (31). Potential short linear motifs (SLiMs) within the TAD were retrieved from the Eukaryotic Linear Motif (ELM) resource database (32).

### Plasmid generation, recombinant protein expression, and purification

A mammalian pcDNA3.1/HisC expression vector harboring the cDNA sequence of full-length *Hs*HNF-1A (33) was used to generate a bacterial expression construct for DBD-TAD (residues 83-631) by Gateway® cloning technology (34). Two PCR reactions were conducted in order to introduce an N-terminal Tobacco-Etch Virus protease site (ENLYFQG) and attB1/2 sites required for recombination. A BP reaction was performed to transfer the gene sequence into the pDONR221 entry vector (Invitrogen). A subsequent LR reaction was conducted for DNA insert transfer into the pTH27 (35) destination vector, generating a bacterial expression plasmid encoding for a His_6_-TEV-DBD-TAD construct (hereafter DBD-TAD). Sequence identity and integrity were verified by plasmid DNA sequencing. Primers used in cloning and sequencing are listed in Table 1.

**Table 1.**
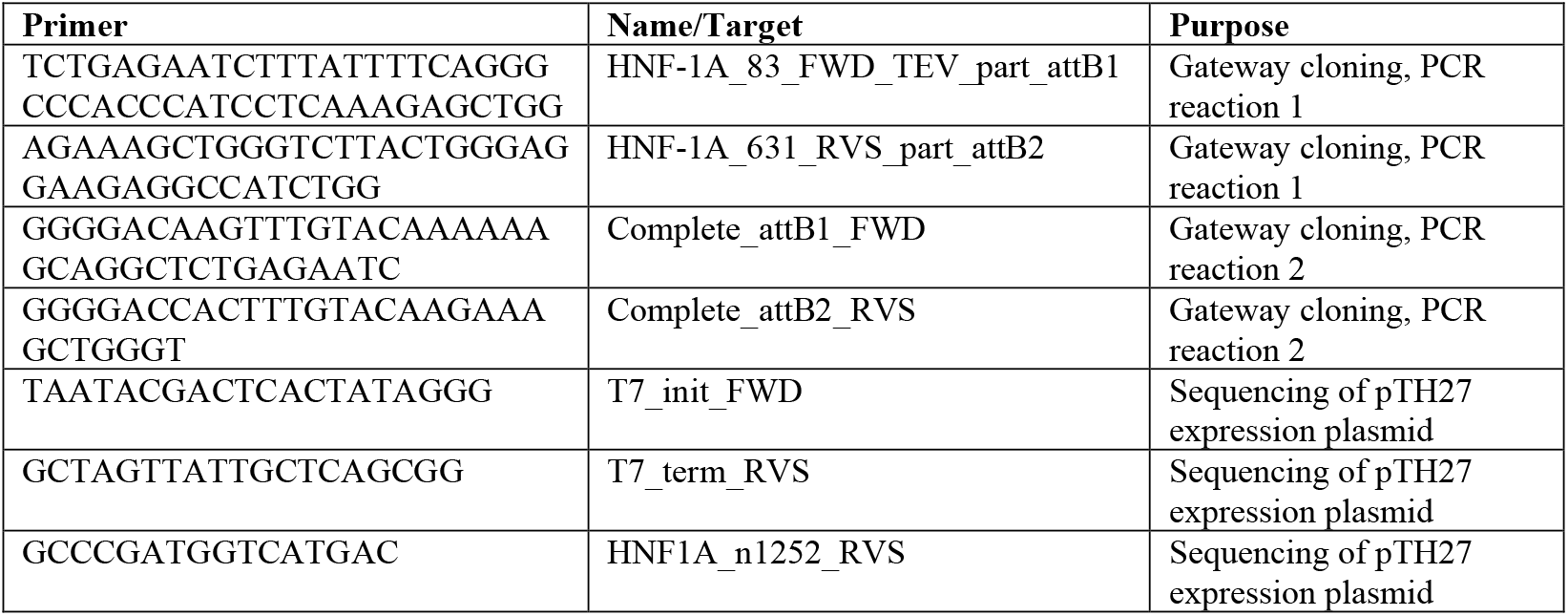
Primers used for the generation and sequencing of the His^6^-TEV-DBD-TAD expression plasmid.

DBD-TAD was recombinantly expressed in *E. coli* Rosetta(DE3). Bacteria were cultured in LB medium at 37 °C until an OD_600_ value of 0.6-0.8 was reached. Protein expression was induced by the addition of 1 mM IPTG and conducted at 18 °C for 20 h. Bacteria were harvested by centrifugation (6 000 × *g*, 10 min, 4 °C) and resuspended in lysis buffer containing urea as denaturation reagent (50 mM Tris pH 8.5, 500 mM NaCl, 6 M urea, 10 mM imidazole, 1 mM DTT, 1 mM PMSF, 1× cOmplete EDTA-free protease inhibitor cocktail). The sample was ultrasonicated (7 min, 1 s on/off cycles, 25 W), and the lysate was cleared by centrifugation (16 000 × *g*, 30 min, 4 °C).

The soluble fraction was loaded onto a pre-equilibrated Ni-NTA column with a 2 ml bed volume. The column was washed with 15 column volumes (CVs) of wash buffer (50 mM Tris pH 8.5, 500 mM NaCl, 6 M urea, 10 mM imidazole, 1 mM DTT, 1 mM PMSF), before elution was performed using a stepwise imidazole gradient (40 mM, 60 mM, 80 mM, 300 mM imidazole) and a fraction volume of 3 CVs. Fractions containing the protein of interest were pooled and dialyzed into a buffer lacking urea (20 mM Tris pH 8.5, 500 mM NaCl, 1 mM DTT), using a dialysis tubing with a 12-14 kDa molecular weight cut-off (Spectra/Por). The dialyzed sample, containing the refolded His-tagged DBD-TAD protein, was concentrated and subjected to size exclusion chromatography (SEC). A Superdex 200 10/300GL Increase column (GE Healthcare) was used at a flow rate of 0.5 ml/min with a running buffer containing 20 mM Tris pH 8.5, 500 mM NaCl, 1 mM TCEP. Pooled SEC fractions containing His-tagged DBD-TAD were concentrated, snap-frozen in liquid N_2_, and stored at -80 °C.

The DBD of HNF-1A (residues 83-279) was recombinantly expressed and purified as previously described (12) and used for control purposes.

### Synchrotron radiation circular dichroism

SRCD experiments were performed on the AU-CD beamline, ASTRID2 synchrotron (Aarhus, Denmark). DBD and DBD-TAD were dialyzed into SRCD buffer (20 mM sodium phosphate pH 7.8, 150 mM NaF, 1 mM TCEP) prior to measurements. Samples with various 2,2,2-trifluoroethanol (TFE) concentrations were prepared by mixing with 100% TFE (Sigma-Aldrich T63002). All samples were measured at an HNF-1A concentration of 0.5 mg/ml, except for DBD-TAD in the presence of 70% TFE, where the HNF-1A concentration was 0.45 mg/ml. SRCD measurements were performed at 10 °C, using 0.1 mm quartz cuvettes (Hellma Analytics). Three wavelength scans per sample were recorded, applying a scan range of 170-280 nm with a 1 nm step size. Data processing was done using Microsoft Excel and CDtoolX (36).

### Small-angle X-ray scattering

SAXS experiments were performed on the CoSAXS beamline (37), MAX IV Laboratory (Lund, Sweden). DBD-TAD was measured at ∼2.5 mg/ml in batch mode (4 mM Tris pH 8.5, 100 mM NaCl, 1 mM TCEP). Measurements were performed at 10 °C at a wavelength of 1 Å (12.4 keV). Data were collected over an angular range of 3.8 × 10^−4^ – 6.4 × 10^−1^ Å^-1^ with 300 frames at an exposure time of 20 ms per frame. Final sample and buffer scattering curves were obtained by averaging the frames that matched each other to avoid incorporation of radiation damage.

Data reduction and analysis were done using the ATSAS 3.1.0 package (38). Guinier analysis in PRIMUS (38) (range 25 – 38; 1.6 × 10^−2^ – 2.4 × 10^−2^ Å^-1^) yielded an *R*_*g*_ value of 48.5 Å and *I*_*0*_ of 0.039 with a fidelity of 0.72. The scattering data were also analyzed using the particle distance distribution function p(r) generated by GNOM (39) (range 15 – 239; 9.2 × 10^−3^ Å^-1^ – 1.5 × 10^−1^ Å^-1^). The data fit with the highest quality (fidelity = 0.79) was generated when *D*_*max*_ was 170 Å. The results from GNOM agreed with Guinier analysis (*R*_*g*_ = 48.2 Å, *I*_*0*_ = 0.04). The *R*_*g*_ was determined to 51.0 Å by using the Debye formalism essentially as described in (40) and according to original methodology (41-43). Molecular weight analysis in PRIMUS resulted in differing molecular weight estimates depending on the algorithm used (Q_p_: 82.7 kDa, MoW: 35.0 kDa, V_c_: 52.8 kDa, Size and Shape: 126.8 kDa, Bayesian Inference: 91.2 kDa with a credibility interval of [49.3 kDa, 134.3 kDa] at 90.3% probability). The molecular weight estimate based on absolute scale was 55.4 kDa, which was relatively close to the theoretical molecular weight of the His_6_-DBD-TAD protein (62.2 kDa).

The ensemble optimization method (EOM) algorithm was used to generate an ensemble of models satisfying the SAXS data (44, 45). The crystal structure of the DBD (PDB: I1C8, (14)) was used to provide rigid bodies for POU_S_ (residues 87-180) and POU_H_ (residues 209-276). The EOM algorithm was run for native-like behavior of disordered regions, with the POU_S_ domain being fixed in position. The EOM conformers were visualized using PyMOL (46).

### Differential interference contrast microscopy

Purified DBD-TAD and DBD were diluted in storage buffer (20 mM Tris pH 8.5, 500 mM NaCl, 1 mM TCEP), yielding a final concentration of 10 – 25 µM. PEG8000 was added to a final concentration of 10% w/v. For each protein concentration, a 10 µl sample was prepared on a microscopy slide and differential interference contrast (DIC) images were acquired using a Zeiss Axiovert 200M wide-field fluorescence microscope equipped with a LD Plan-NEOFLUAR 40x/0.6 Ph2 objective and an AxioCam HR camera. Image acquisition was performed using the AxioVision software (Carl Zeiss, version 4.5).

### Immunofluorescence microscopy

MIN6 cells (47, 48), kindly provided by Professor Claes Wollheim (Lund University, Sweden), were cultured in DMEM growth medium (Gibco), supplemented with 15% fetal bovine serum and 1% v/v penicillin-streptomycin (Sigma-Aldrich P4333). Cells were incubated at 37 °C and 5% atmospheric CO_2_ in a humidity incubator. For immunofluorescence (IF), MIN6 cells were seeded on coverslips in 24-well plates. IF staining was performed at room temperature (RT). 24 h post-seeding, cells were washed twice with phosphate-buffered saline (PBS) and fixed by incubation in a phosphate buffer containing 3% paraformaldehyde (Sigma-Aldrich) for 25 min. Samples were washed three times with PBS, permeabilized in 0.1% Triton X-100 (Sigma-Aldrich) in PBS for 10 min, and washed three times with PBS. Binding epitopes were blocked by incubation with 2% normal goat serum (GS, Invitrogen)/8% bovine serum albumin (BSA, Sigma-Aldrich) in PBS for 1 h, followed by three PBS wash steps. Primary anti-HNF-1A antibody (Invitrogen, PA5-83263) was applied for 1 h, using a 1:100 dilution in 2% GS/8% BSA-PBS. Samples were washed three times and stored in PBS overnight at 4 °C. Secondary antibody (Alexa Fluor® 488 AffiniPure Goat Anti-Rabbit IgG (H+L), Jackson ImmunoResearch, 111-545-003) was applied for 45 min, using a 1:100 dilution in 2% GS/8% BSA-PBS. Samples were washed three times with PBS and mounted on glass slides using ProLong™ Diamond Antifade DAPI mountant (Invitrogen, P36962). Mounting solution was allowed to solidify overnight, and the slides were inspected on the following day. Samples were imaged using a Zeiss Axiovert 200M wide-field fluorescence microscope equipped with an AxioCam HR camera, a Plan-NEOFLUAR 100X/1.30 Ph3 oil immersion objective and filters appropriate for the detection of DAPI and GFP/Alexa-488 signals. Images were acquired by using the AxioVision software (Carl Zeiss, version 4.5) and superposed using the Fiji ImageJ software (Wayne Rasband, National Institutes of Health, USA, version 1.52a) (49).

### Droplet simulation protocol

Molecular dynamics simulations of the LLPS behavior of HNF-1A TAD (residues 280-631) were performed using the 1 bead per amino acid (1BPA) molecular dynamics model (50, 51), with updated parameters described in (52). The initial cubic simulation box is populated with molecules (using a random initial conformation) with their center of mass placed upon a regular grid, with a small buffer region to avoid overlap between molecules. All simulations are carried out at a temperature of 300 K, 150 mM ion concentration (κ = 1.27 nm^-1^), and use a timestep of 20 fs. For equilibration of the droplet, energy minimization on the initial configuration is used (energy tolerance of 1 kJ mol^-1^ nm^-1^), before 50 ns NVT Langevin dynamics simulations (Nosé-hoover thermostat with τ_t_ = 100 ps), followed by 500 ns NPT Langevin dynamics (Nosé-hoover thermostat with τ_t_ = 100 ps and a Berendsen barostat with τ_p_ = 10 ps, 1 bar reference pressure and a compressibility of 4.5 10^−5^ bar^-1^). The end state of the NPT equilibration step is inserted into a new periodic box with a volume chosen to give a total residue density of 80 mM, after recentering on the center of mass and after the molecules have been unwrapped across the previous periodic boundary conditions. A second energy minimisation step is applied in the new simulation box to relax the molecules after the box expansion (energy tolerance of 1 kJ mol^-1^ nm^-1^). A final 3 μs NVT equilibration/production run (Nosé-hoover thermostat with τ_t_ = 100 ps) is used for data collection. The trajectory is sampled every 5 ns to determine whether convergence was reached.

## Results and Discussion

### Structure prediction of HNF-1A

Bioinformatic sequence analyses provided insights into structural features and sequence characteristics of human HNF-1A (Fig. 1). Disorder prediction using IUPred3 was performed to analyze the residue-dependent propensity of HNF-1A to fold into globular domains or remain disordered (Fig. 1A) (23). While the α-helical DD and DBD of HNF-1A were predicted to undergo protein folding, the linker between DD and DBD showed a high tendency to remain disordered (Fig. 1A). The flexibility of this linker region has previously been demonstrated by SAXS (12), illustrating the agreement between *in vitro* experiments and *in silico* prediction. The IUPred3 prediction indicated that the TAD likely contains long IDRs. Disorder was predicted for the residue ranges 280-420, 460-500, and 540-590, while the regions harboring residues 420-460, 500-540, and 590-631 may fold in a context-specific manner (Fig. 1A). A complementary analysis using the ANCHOR2 algorithm was in agreement with the IUPred3 prediction (Fig. 1A). The residue ranges 300-400 and 550-600 were predicted to contain disordered binding sites that can undergo disorder-to-order transitions upon binding to a partner protein (Fig. 1A).

Tertiary structure predictions of HNF-1A using AlphaFold (26, 27) were in accordance with IUPred3 and ANCHOR2 (Fig. 1B). The per-residue estimate of prediction confidence (pLDDT score) was <50 for residues 300-631, indicating a disordered conformation for this protein region. Apart from a C-terminal extension of helix α8 in the DBD and several low-confidence short helices in the TAD, the AlphaFold prediction suggested an entirely disordered TAD (Fig. 1B). However, context-dependent structural changes predicted by the ANCHOR2 algorithm (Fig. 1A) may not be represented in this Alphafold model. Such folding events, which follow a coupled folding and binding mechanism, have been demonstrated for IDRs in other transcription factors (e.g. DREB2A, p53, HIF1α) when binding to specific interaction partners (19, 53, 54).

### Sequence composition of the TAD

A general feature of intrinsically disordered proteins (IDPs) is the absence of hydrophobic clusters and an enrichment of charged and polar residues (55). Indeed, a hydrophobicity analysis using FELLS illustrated the lack of hydrophobic residues in the TAD compared to the globular DD and DBD in the N-terminal half of HNF-1A (Fig. 1C). Interestingly, charge distribution analysis of the HNF-1A sequence revealed that the TAD contains very few charged residues (Fig. 1C). This observation was surprising, as charged amino acids generally promote solubility and are frequently found in IDPs (55). However, the sequence analysis also revealed that the TAD is highly enriched in Ser and Thr residues. Such a polar tract may facilitate the interaction with the polar environment and promote a disordered state of the TAD (Fig. 1C).

In addition to modulating protein solubility, the side chains of Ser and Thr residues can be phosphorylated, leading to changes in protein function (56). The high prevalence of Ser and Thr residues in the TAD of HNF-1A may indicate that it is a target of post-translational modification by kinases and phosphatases. As a transcription factor, HNF-1A may be phosphorylated by kinases within the basal transcriptional machinery, e.g. cdk7-9, TAFII250, or TFIIF (57). Cdk7 phosphorylates the activation domains of the transcription factors ERα, E2F-1, and p53, which results in changes in their transactivating properties (57). Such phosphorylation events may dynamically affect the ability of HNF-1A to interact with transcriptional co-activators/repressors and chromatin remodeling proteins, modulate the localization of HNF-1A, or mediate changes in protein turnover.

The most common activation domains in transcription factors are acidic activation domains, which require a balance between acidic and hydrophobic residues (58, 59). PADDLE, a recently developed algorithm to identify such domains in protein sequences (59), did not predict any acidic activation domains within the TAD of HNF-1A (data not shown). The transactivation potential of the TAD may thus unfold via a different mechanism. Considering the sequence composition of the TAD (Fig. 1C), we speculate that activation domains rich in Gln, Ser, and Pro may play a role in the function of the HNF-1A TAD (60, 61).

In recent years, researchers have uncovered the ability of transcription factors to activate gene transcription by undergoing LLPS. The formation of transcriptional condensates is often mediated via activation domains and leads to the dynamic compartmentalization of the gene transcriptional machinery, including RNA polymerase II and its cofactors (62, 63). We hypothesized that the HNF-1A TAD may have the potential to activate gene transcription in the same way. We utilized the FuzDrop algorithm to compute the probability of HNF-1A to undergo spontaneous LLPS (28), and the obtained pLLPS score of 0.9932 suggested a strong LLPS tendency. The computed droplet-promoting regions (DPRs) were found across the entire HNF-1A sequence (Fig. 1D). Interestingly, some of the predicted DPRs overlapped with the previously identified activation domains in the TAD, which may hint towards a transactivation mechanism involving LLPS. FuzDrop also predicted HNF-1A to contain aggregation hotspots, which partially overlapped with the predicted DPRs (Fig. 1D).

### Sequence conservation and short linear motifs in the TAD

We generated a multiple sequence alignment to investigate the sequence conservation of the TAD (Fig. 1E). Some TAD regions are highly conserved across species, such as the residue ranges 374-387, 457-469, and 487-496, while conservation in other regions is mainly restricted to mammals, e.g. in residue ranges 285-311, 330-350, and 571-589.

IDRs frequently harbor SLiMs, which are sequence elements of 6 -12 residues that mediate the binding to folded interaction partners or promote LLPS by partaking in multivalent interactions (64). In order to identify potential SLiMs in the TAD of HNF-1A, we performed a search in the ELM resource (32). High-confidence SLiMs are marked in the multiple sequence alignment (Fig. 1E) and include interaction sites for WW domains (residues 287-308, residues 555-560), SH3 domains (residues 287-308), FHA domains (residues 525-530), as well as a recognition site for USP7 ubiquitination enzymes (residues 626-631). Interestingly, many of these SLiMs overlapped with recognition sites for different kinases, such as MAPK, Cks1, and NEK2, which may promote Ser or Thr phosphorylation within the motifs and thereby alter their ability to interact with the respective protein domains. The predicted SLiMs were localized in both highly conserved and less conserved regions (Fig. 1E).

In conclusion, our sequence analyses provided a useful starting point for the functional dissection of the HNF-1A TAD. Experimental studies will be required in the future to investigate the biological roles of specific TAD regions and their dynamic regulation. Biochemical and structural approaches will shed light on the structure-function relationships of HNF-1A in solution.

### Recombinant expression and purification of a TAD-containing HNF-1A protein

In order to provide a basis for functional studies, we set out to biophysically and structurally characterize the TAD of HNF-1A. We generated three N-terminally His_6_-tagged HNF-1A expression constructs harboring the TAD (DD-DBD-TAD, residues 1 – 631; DBD-TAD, residues 83 – 631; TAD, residues 280 – 631). Initial overexpression in Rosetta(DE3) resulted in insoluble protein, which may be functionally connected to the predicted LLPS or aggregation behavior of the protein (Fig. 1D). We screened for optimal expression and lysis conditions by varying expression systems (Lemo21(DE3) *E. coli*, Sf9 *Spodoptera frugiperda*), expression conditions (1 – 72 h, 20/37 °C), and lysis buffer compositions (screening for NaCl concentrations, pH value, and stabilizing additives). However, none of these adjustments yielded soluble protein (data not shown).

We proceeded with protein purification from the insoluble lysate fraction, whereby we focused on the DBD-TAD construct due to relatively high expression levels. We solubilized the protein under denaturing conditions during bacterial lysis and performed a first purification step using Ni-NTA affinity chromatography. The protein did not have a strong affinity towards the Ni-NTA material, as relatively low imidazole concentrations (40 – 80 mM) were sufficient for elution. Moreover, the elution fraction contained three strong bands, indicating a C-terminal truncation of overexpressed DBD-TAD. In order to refold the denatured protein into its native form, we dialyzed the elution fractions into buffer lacking denaturant. We performed a final size exclusion chromatography (SEC) step to separate monomeric DBD-TAD from aggregates and suspected degradation products (Fig. 2A, B). Mass spectrometry confirmed that both the purified protein and the suspected degradation products were derived from human HNF-1A. As the purified protein had shown a tendency for C-terminal degradation, we aimed to conduct all characterization experiments immediately after purification.

**Fig. 2.**
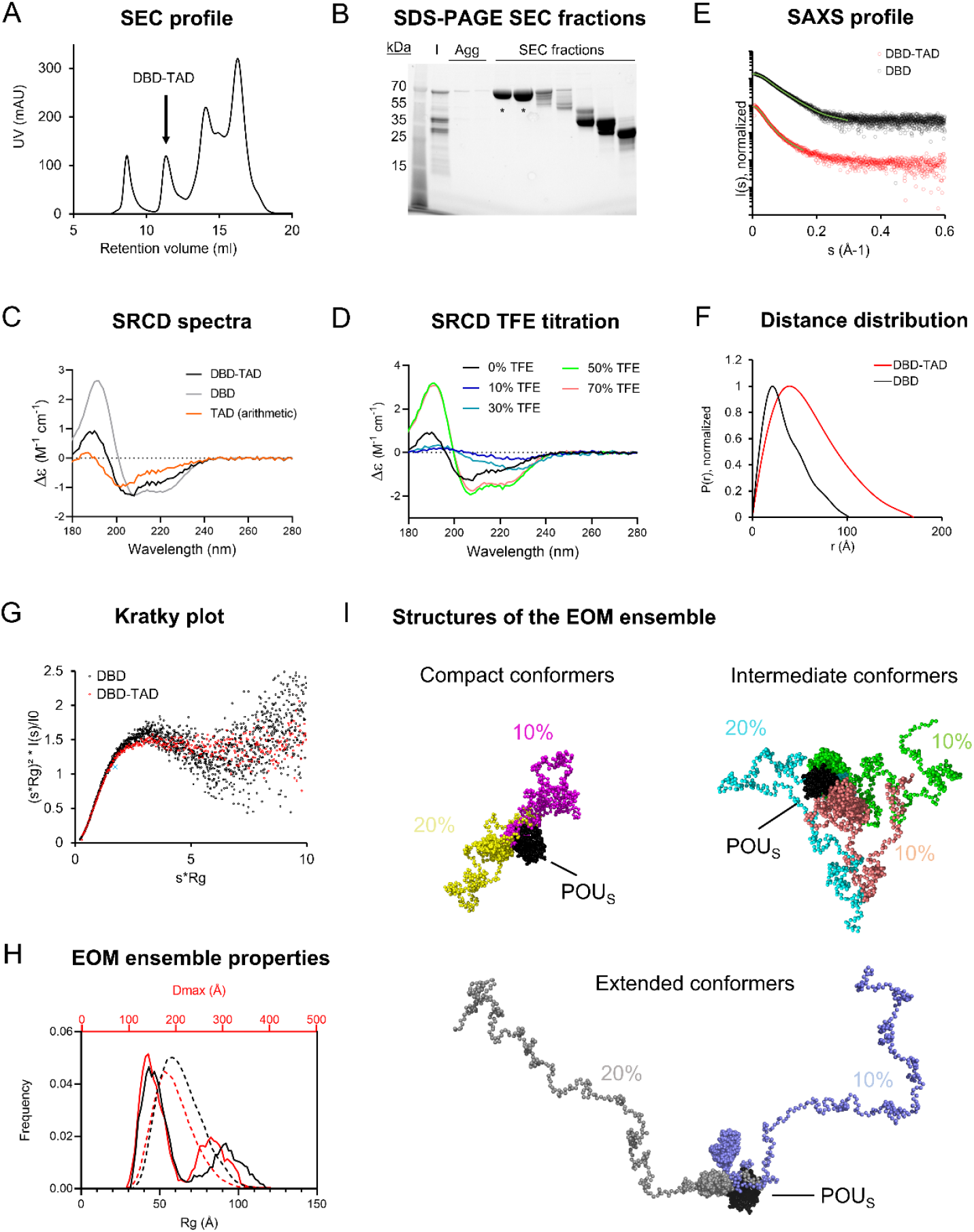
Purification and biophysical characterization of the DBD-TAD construct. **A**. SEC profile with the DBD-TAD containing peak indicated. **B**. SDS-PAGE analysis of respective SEC fractions from various peaks of the chromatogram shown in A. Agg -SEC fractions containing aggregated proteins. The bands marked with ‘*’ were pooled and used for experiments. Lower molecular weight species in the residual SEC fractions corresponded to degradation products of the recombinant protein. **C**. SRCD spectra for DBD (grey) and DBD-TAD (black). A theoretical spectrum for the isolated TAD (orange) was calculated by subtraction of the two obtained spectra. **D**. TFE titration experiment for DBD-TAD. **E-G**. SAXS data analyses for DBD-TAD. DBD SAXS data, previously published by us (12), are presented for direct comparison. **E**. Scattering curve. **F**. Distance distribution function. **G**. Normalized Kratky plot, with the cross indicating the expected maximum for a globular particle (v3, 1.104) (65). **H**. *D*^*max*^ (red) and *R*^*g*^ (black) distributions from EOM analysis for DBD-TAD, illustrating ensemble frequencies as solid lines and pool frequencies as dashed lines. **I**. Seven conformer structures in the generated EOM ensemble with assigned abundance values, each represented in a different color. The models were superposed based on the POU^S^ domain (black). Conformers are grouped by the degree of compaction.

### The HNF-1A TAD is intrinsically disordered as revealed by SRCD and SAXS

SRCD experiments were performed to assess the secondary structure content of the purified protein (Fig. 2C). The SRCD spectra indicated that DBD-TAD contained both α-helical structures and random-coil regions. Since the isolated DBD produced an SRCD spectrum indicative of α-helical structure content (Fig. 2C), we concluded that the TAD of HNF-1A likely adopts a random-coil structure in solution. We performed a subtraction of the DBD-TAD and DBD spectra, yielding a theoretical TAD spectrum, which indicated a disordered protein conformation and confirmed our hypothesis. To test the propensity of the TAD to fold into an α-helical structure, we performed a TFE titration experiment, in which α-helical structure of the DBD-TAD construct was induced by the addition of ≥50% TFE (Fig. 2D). Notably, intermediate TFE concentrations (10-30%) decreased the SRCD signal across the entire wavelength range, which may be due to secondary processes causing light scattering, such as LLPS or aggregation.

We employed SAXS to study the molecular dimensions and shape of DBD-TAD (Fig. 2E-G). We compared the scattering data to a published SAXS dataset for the isolated DBD (12), allowing us to study the contributions of the HNF-1A TAD. The DBD-TAD sample was free of aggregation (Fig. 2E). The molecular dimensions of the protein were determined using the distance distribution function (*R*_*g*_ = 4.8 nm, *D*_*max*_ = 17 nm). DBD-TAD presented larger *R*_*g*_ and *D*_*max*_ compared to the isolated DBD (*R*_*g*_ = 2.8 nm, *D*_*max*_ = 10 nm (12)), which was expected due to the higher molecular weight of DBD-TAD. The dimensionless Kratky plot indicated that DBD-TAD was highly flexible and had a non-globular shape (Fig. 2G). As this behavior was more pronounced than for the isolated DBD protein, we attributed this protein property to the disordered TAD (Fig. 2G). Based on the scattering data, we generated a SAXS model to visualize the molecular shape of the protein (Fig. 2H-J). We utilized EOM to generate an ensemble of conformers that together satisfy the SAXS data. A published crystal structure of a DBD:DNA complex (PDB: 1IC8 (14)) was used to model POU_S_ and POU_H_ as globular domains. Missing residues were modeled as flexible regions. The data were satisfied by an ensemble containing seven conformers with approximately equal abundance, exhibiting an average *R*_*g*_ value of 64 Å and a *D*_*max*_ value of 198 Å. An analysis of the ensemble properties revealed a bi-modal distribution of the *R*_*g*_ and *D*_*max*_ parameters (Fig. 2H), with one population exhibiting a peak *R*_*g*_ of ∼45 Å and a peak *D*_*max*_ of ∼140 Å, and another population exhibiting higher molecular dimensions with a peak *R*_*g*_ of ∼90 Å and a peak *D*_*max*_ of ∼280 Å. This is reflected in the generated conformer models, in which the TAD is present in compact, intermediate, and extended conformations (Fig. 2I).

### The TAD promotes LLPS of HNF-1A *in vitro*

Based on the FuzDrop predictions (Fig. 1D), we suspected that the disordered TAD may have the potential to promote LLPS of HNF-1A. We observed that the purified DBD-TAD protein could not be concentrated above ca. 2.5 mg/ml. The protein solution increased in viscosity at higher concentrations, which may be due to the LLPS-driven formation of protein droplets, hydrogel or fibrous aggregates (66). To investigate the potential cause, we performed DIC microscopy using purified DBD-TAD and DBD (Fig. 3A, B). We used the crowding agent PEG8000 to mimic the dense molecular environment of the nucleus. We observed droplet formation for the DBD-TAD construct, with a concentration-dependent increase in droplet number (Fig. 3A). Most droplets had a round shape, suggesting that they were liquid condensates, as opposed to aggregates. The DBD protein also produced droplets; however, higher protein concentrations were required, and the droplet size differed compared to the TAD-containing protein (Fig. 3B). The latter indicates that the TAD may be a major driver of the LLPS behavior of HNF-1A. However, it is important to note that the comparison between the two proteins and their contribution to LLPS is not straightforward due to their different molecular size. Additional experimental evidence for the presence of HNF-1A-containing droplets *in vitro* may be acquired by performing fluorescence microscopy with a fluorescently labeled (DBD-)TAD, observing the mixing dynamics and potential fusion events between the droplets (67, 68).

**Fig. 3.**
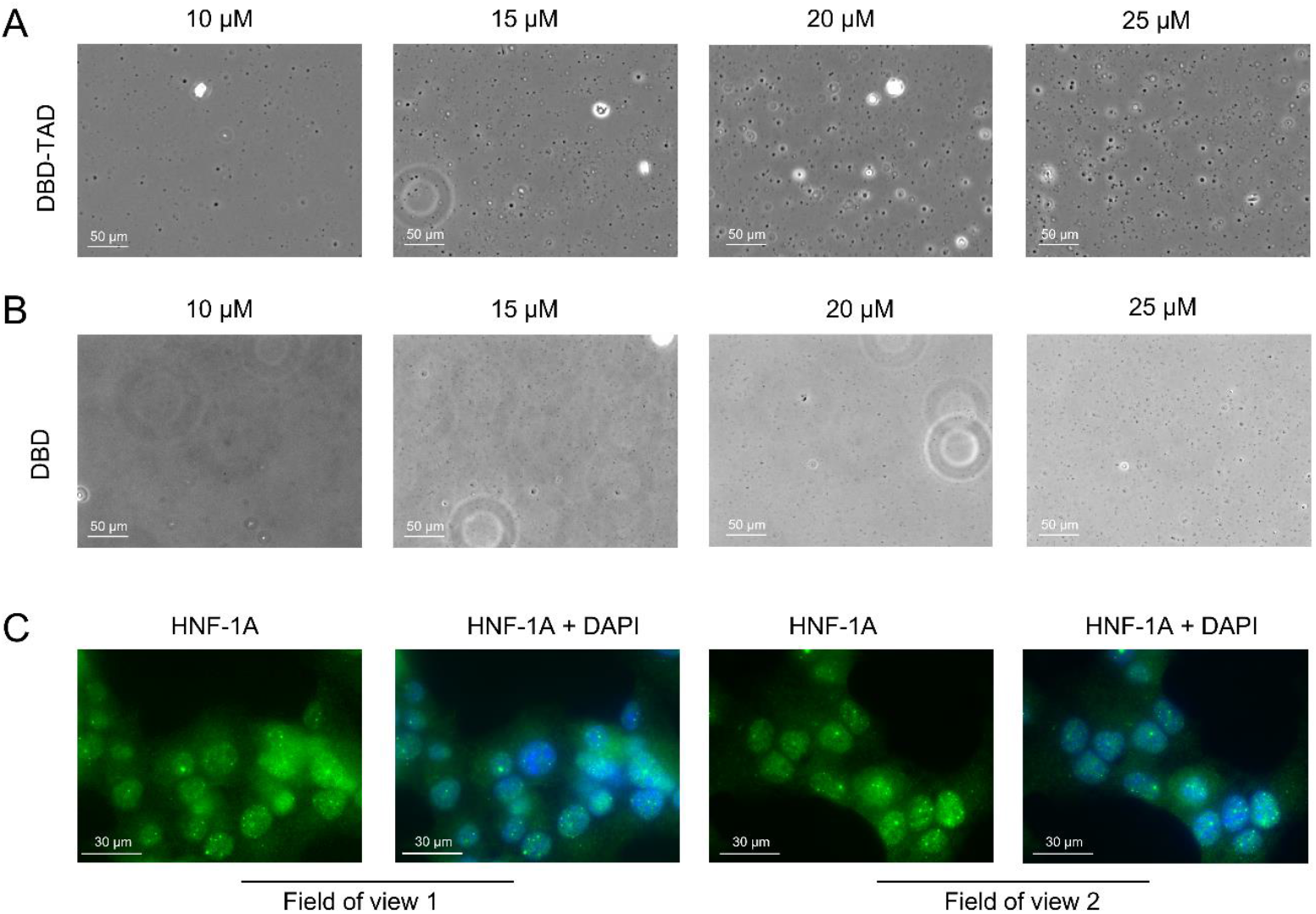
Initial evidence for LLPS behavior of HNF-1A. **A-B**. Representative DIC microscopy images of purified DBD-TAD (**A**) or DBD (**B**) at different concentrations (10 – 25 µM) in the presence of 10% PEG8000 as a molecular crowding agent. **C**. IF images of MIN6 β-like cells, stained for HNF-1A using an HNF-1A specific antibody and DNA using DAPI stain. Two fields of view from the same microscopy slide are presented, where the left image represents HNF-1A signals (green) and the right image a superposition of the same HNF-1A signal (green) and nuclear DAPI signal (blue).

As transcription factors are normally present in the nM concentration range (copy numbers between 10^3^-10^6^ per mammalian cell) (69, 70), our experimental conditions do not fully recapitulate the protein concentration at endogenous levels. However, even with lower HNF-1A concentrations in the cell, phase separation may take place in the presence of other IDR-containing proteins promoting multivalent interactions. LLPS-dependent cellular compartments typically concentrate ten to several hundred different proteins (and often RNA molecules), whereby many of them contribute to the formation of the biomolecular condensate (71). We next investigated whether the suspected LLPS behavior of HNF-1A may lead to the formation of biological condensates in cells. We employed fluorescence microscopy to image the localization of endogenous full-length HNF-1A in fixed MIN6 β-like cells, revealing that HNF-1A was predominantly located in the nucleus (Fig. 3C). Moreover, HNF-1A produced strong fluorescent punctae, which co-localized with the nuclear stain and may thus represent nuclear condensates (Fig. 3C). It is important to note that the cell fixation using paraformaldehyde can significantly alter the appearance or presence of biomolecular condensates, which is why the presented results should be interpreted with care and verified with alternative methods (72). Live-cell imaging techniques will be valuable for verifying the existence of HNF-1A containing nuclear condensates, and further molecular biological studies will help to elucidate their components and biological function.

### TAD undergoes LLPS in molecular dynamics simulations

To understand the LLPS propensity of the HNF-1A TAD, we also undertook computational droplet formation simulations (Fig. 4, S1, Video V1). We assessed the internal structure of the droplets by using the 1 bead per amino acid (1BPA) molecular dynamics model (52), developed for the study of intrinsically disordered proteins in the nuclear pore complex (50, 51). Self-assembly and clustering of individual monomers into phase-separated condensates can be a slow process to observe. To speed up this process, a condensed phase droplet is formed at the start of the simulation, which is then inserted into an empty dilute phase. If LLPS is favored, the droplet structure should remain stable throughout the subsequent simulation; if non-favored, the droplet would break up into a dilute phase of monomers.

**Fig. 4.**
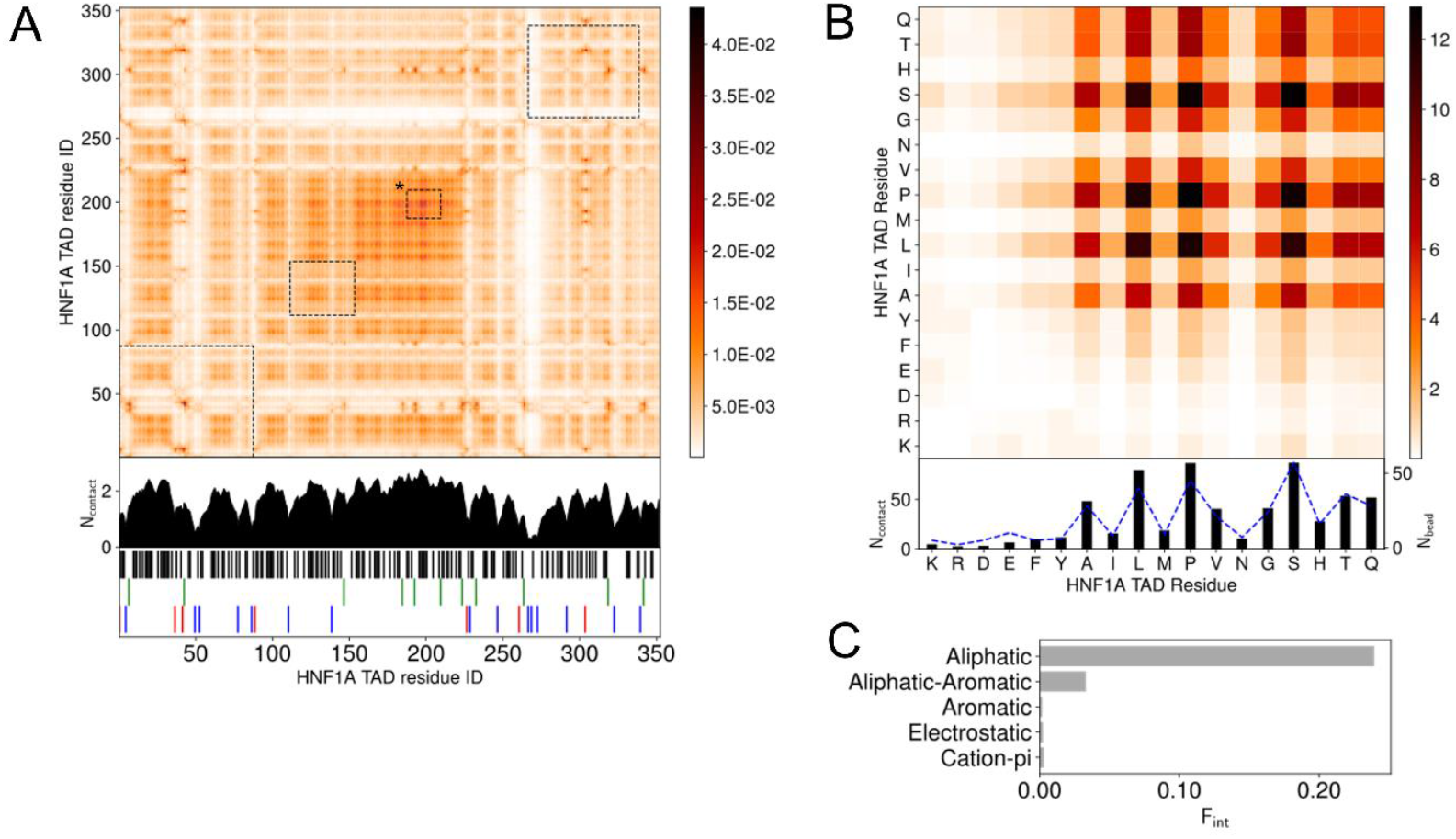
LLPS promoting TAD interactions, studied by droplet simulations. **A**. Intermolecular contact map by residue index for 120 HNF-1A TAD molecules at 150 mM ion concentration and 300 K. The contacts in the contact maps by residue index are averaged in time (600 frames) and normalized by the number of HNF-1A TAD molecules (120) in the simulation. The 1D summation is shown below the contact map for each residue. Residues are categorized into 5 groups: cations (R, K)-red, anions (D, E)-blue, aromatic (F, Y, W)-green, aliphatic (A, C, I, L, M, P, V)-black, hydrophilic (G, N, S, H, Q, T)-white. The interactions between droplet promoting regions predicted by FuzDrop (Fig. 1D) are shown by black dashed boxes. Box marked with ‘*’ corresponds to FuzDrop DPR 466-488. **B**. Intermolecular contact map by residue type for 120 HNF-1A TAD molecules. The contact map is a matrix reduction of the contact map by residue index in (A). The abundance for the residues, N^bead^, are shown by blue dashed lines. **C**. Interaction summary for a droplet simulation with 120 HNF-1A TAD molecules at 150 mM ion concentration and 300 K. The fraction of interactions, F^int^, are aggregated by type and normalized by the total number of interactions.

We simulated the behavior of 120 HNF-1A TAD molecules and observed LLPS (Video V1). In the beginning of the simulation, two to three small clusters with one to six HNF-1A TAD molecules co-existed with a large cluster containing approximately 115 HNF-1A TAD molecules. The smaller clusters rapidly fused with the big cluster, yielding a droplet of 120 HNF-1A TAD molecules that remained stably associated throughout the simulation period (Video V1). By analyzing the interactions between the TAD molecules in detail, we found that intermolecular interactions along the whole length of the TAD contribute to LLPS (Fig. 4A). The LLPS behavior of the TAD is driven by hydrophobic aliphatic-aliphatic contacts, with minimal aliphatic-aromatic interactions due to the low abundance of aromatic residues (Fig. 4C). This is driven by contacts between Ala, Leu, Pro, and Val (Fig. 4B). We also observed many interactions between hydrophilic residues (Gly, Ser, His, Thr and Gln) driven by their high abundance (Fig. 4B), but whose effect is reduced when interactions are normalized by amino acid abundance (Fig. S1). The predicted LLPS-promoting regions (Fig. 1D) are highlighted on the contact map in Fig. 4A. These regions partially align with the regions we found to form the most contacts, with only the FuzDrop DPR 466-488 showing full alignment (box marked with ‘*’ in Fig. 4A). The more extensive interactions discovered by the simulations indicate that the even distribution of aliphatic residues throughout the TAD contributes to LLPS, whereas an absence of intermolecular interactions among charged and polar residues, which make up 90% of the TAD, was evident (white stripes in Fig. 4A). In summary, the results of our simulations corroborate with our *in vitro* data (Fig. 3A) and suggest that several residues along the entire TAD collectively contribute to the droplet formation.

## Conclusions

Our study provides insights into the sequence features and structural properties of the uncharacterized TAD of HNF-1A. We developed the first purification protocol for a TAD containing HNF-1A construct, which allowed us to study the TAD *in vitro*. We found that the TAD is intrinsically disordered, which may be crucial for the dynamic interaction with other proteins involved in gene transcriptional control. While we reproducibly obtained pure DBD-TAD samples, we observed a tendency for protein degradation from the C-terminal end. Further efforts in construct design and protein purification may thus be undertaken to improve the yield and protein stability to allow for an improved structural analysis of the TAD. Our functional data on the potential LLPS behavior of HNF-1A agreed with predictions and *in silico* simulations and supports the hypothesis that the TAD may drive the formation of transcriptional condensates. Our study provides a foundation for future research on the TADs potential to form condensates and mediate protein-protein interactions, which may be of critical importance for β-cell function. An enhanced understanding of HNF-1A-mediated gene transcription may potentially expose novel treatment targets in patients with HNF1A-MODY.

## Acknowledgements

We are thankful to Ulrich Bergmann (Biocenter Oulu Proteomics Core Facility, University of Oulu, Finland) for mass spectrometry analyses and to Henriette Aksnes (Department of Biomedicine, University of Bergen, Norway) for guidance with fluorescence microscopy. We are grateful for awarded beamtime and thank the support of the synchrotron beamline staff at MAX IV (Lund) and ISA (Aarhus). We thank BioCat - The National graduate school in biocatalysis - for financial support connected to beamtime travel. We acknowledge the use of the Core Facility for Biophysics, Structural Biology, and Screening (BiSS) at the University of Bergen, which has received infrastructure funding from the Research Council of Norway (RCN) through NORCRYST (grant number 245828) and NOR–OPENSCREEN (grant number 245922). This work was funded with a PhD fellowship by the Medical Faculty, University of Bergen, UiB, Norway (to L. K.) and by project grants awarded by the UiB Meltzer foundation (to L. K.). We thank the oLife COFUND project for funding (to M. D. D.). The COFUND project oLife has received funding from the European Union’s Horizon 2020 research and innovation programme under grant agreement No 847675. We thank the Center for Information Technology of the University of Groningen for their support and for providing access to the Peregrine high performance computing cluster. This work made use of the Dutch national e-infrastructure with the support of the SURF Cooperative using grant no. EINF-3233.

