## Supplementary information for "Structural properties of the HNF-1A transactivation domain"

Fig. S1 Molecule contact maps for a droplet simulation with 120 HNF-1A TAD molecules at 150 mM ion concentration and 300 K.

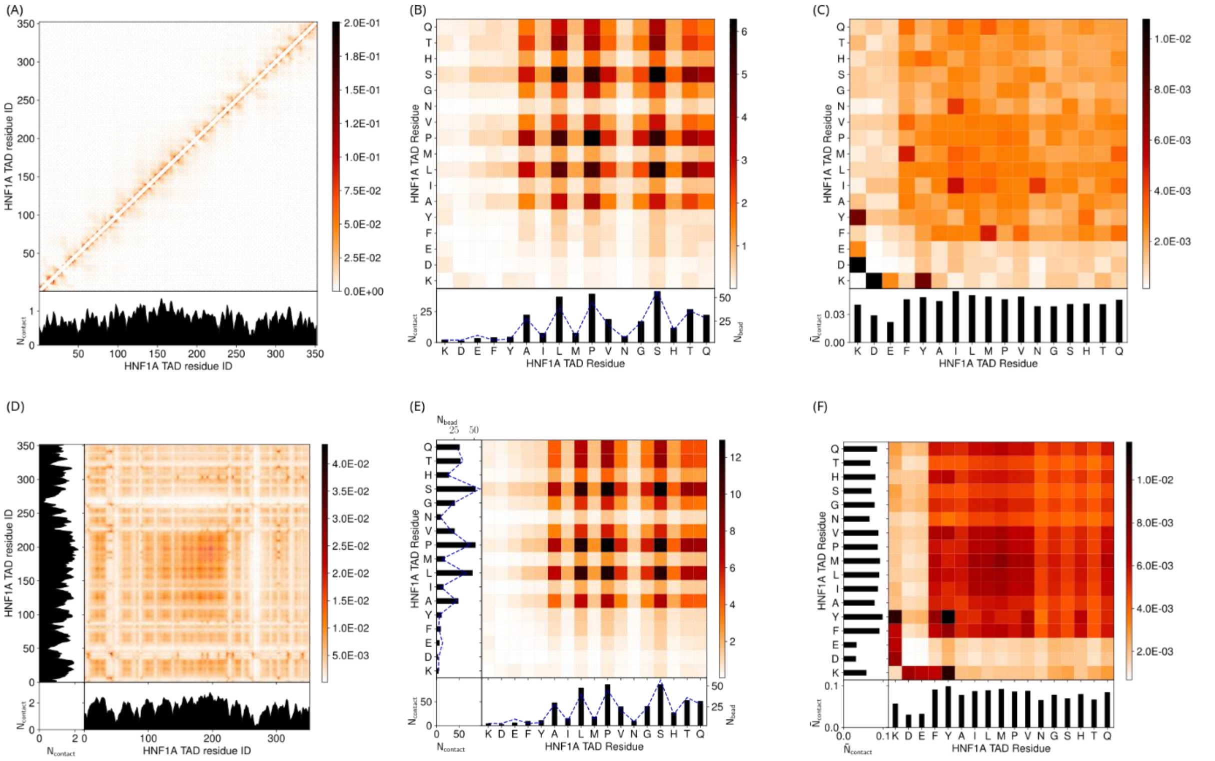


**Figure S1: Single molecule contact maps for a droplet simulation with 120 HNF-1A TAD molecules at 150 mM ion concentration and 300 K.** **A.** Intramolecular contact map by residue index. **B.** Intramolecular contact map by residue type. **C.** Intramolecular contact map by residue type, normalized by residue abundance. **D.** Intermolecular contact map by residue index. **E.** Intermolecular contact map by residue type. **F.** Intermolecular contact map by residue type, normalized by residue abundance. The contacts in the contact maps by residue index (left column) are normalized by the number of frames used (600) and the number of HNF1A TAD molecules (120). The contact maps aggregated by residue type (centre column) are a matrix reduction of the contact map by residue index (left column). The abundance for the residues, $N_{bead}$, are shown by blue dashed lines. The contact maps (right column) are normalized by the square root of the relative bead abundance in the molecules.
